# Gene regulatory logic of the interferon-β enhancer contains multiple selectively deployed modes of transcription factor synergy

**DOI:** 10.1101/2025.02.04.636520

**Authors:** Allison Schiffman, Zhang Cheng, Diana Ourthiague, Alexander Hoffmann

**Affiliations:** Signaling Systems Laboratory, Department of Microbiology, Immunology and Molecular Genetics, and the Institute for Quantitative and Computational Biosciences (QCB), University of California Los Angeles, 611 Charles Young Drive, Los Angeles, CA 90095

**Keywords:** interferon-β, Boolean logic gates, thermodynamic state ensemble models, gene regulatory circuits, NFκB, interferon regulatory factors (IRF), Rig-I, TLR

## Abstract

Type I interferon IFNβ is a key regulator of the immune response, and its dysregulated expression causes disease. The regulation of IFNβ promoter activity has been a touchpoint of mammalian gene control research since the discovery of functional synergy between two stimulus-responsive transcription factors (TFs) nuclear factor kappa B (NFκB) and interferon regulatory factors (IRF). However, subsequent gene knockout studies revealed that this synergy is condition-dependent such that either NFκB or IRF activation can be dispensable, leaving the precise regulatory logic of IFNβ transcription an open question. Here, we developed a series of quantitative enhancer states models of IFNβ expression control and evaluated them with stimulus-response data from TF knockouts. Our analysis confirmed that TF synergy is a hallmark of the regulatory logic but that it need not involve NFκB, as synergy between two adjacent IRF dimers is sufficient. We found that a sigmoidal binding curve at the distal site renders the dual IRF synergy mode ultrasensitive, allowing it only in conditions of high IRF activity upon viral infection. In contrast, the proximal site has high affinity and enables expression in response to bacterial exposure through synergy with NFκB. However, its accessibility is controlled by the competitive repressor p50:p50, which prevents basal IRF levels from synergizing with NFκB, such that NFκB-only stimuli do not activate IFNβ expression. The enhancer states model identifies multiple synergy modes that are accessed differentially in response to different immune threats, enabling a highly stimulus-specific but also versatile regulatory logic for stimulus-specific IFNβ expression.

**SIGNIFICANCE:** Precise regulation of the immune cytokine IFNβ is essential for human health. Classic studies established that the transcription factors NFκB and IRF function synergistically in activating IFNβ expression. However, more recent data revealed that either factor may be dispensable in certain conditions, leaving the regulatory logic of IFNβ transcription an open question. In this study, we evaluated several quantitative models to determine what regulatory logic can account for the available data. We found that the enhancer exhibits multiple synergy modes with specific transcription capabilities that are deployed in a condition-dependent manner. The resulting IFNβ regulatory logic model reveals how stimulus-specificity is achieved, and may be tuned, thereby overcoming a bottleneck in predictive modeling the control of innate immune responses.

## INTRODUCTION

Type I interferons are key regulators of the innate immune response. The most prominent type I interferon, IFNβ, functions both as an autocrine and paracrine cytokine that has important roles in myriad aspects of health and disease. Its primary role is to activate the expression of antiviral genes (1–3) but it also has broad roles in the adaptive immune response by regulating inflammation (4), and the functions of antigen-presenting cells (5, 6), T cells, B cells, and NK cells (7, 8). Conversely, IFNβ dysregulation is associated with diseases (9, 10), as it can lead to systemic activation of the immune response (11, 12), contribute to immunosuppression (2), and impair cell growth and health by downregulating protein synthesis (13, 3). Consequently, IFNβ expression must be tightly tuned to the specific context and trigger stimulus such that it is expressed only when and to the degree needed to overcome the immune threat.

The key regulatory step that controls IFNβ production is the initiation of transcription of the *ifnb1* gene. Its stimulus response is primarily controlled by one κB site, which binds nuclear factor kappa-light-chain-enhancer of activated B cells (NFκB) and two IRE sites, which bind interferon regulatory factors (IRFs). NFκB and IRF are families of stimulus-responsive transcription factors (TFs) which respond to pathogen-associated molecular patterns (PAMPs) via pathogen recognition receptors (PRRs). Indeed, while PRRs are diverse and encompass toll-like receptors (TLRs), Nod-like receptors (NLRs), Rig-I-like receptors (RLRs), and cytosolic DNA sensing receptors (cGAS) (14, 15), they all converge on the activation of NFκB and IRF signaling via their respective MyD88, TRIF, MAVS and STING adaptors (16–19).

Foundational studies of the IFNβ enhancer reported functional synergy between NFκB and IRF, i.e., that more IFNβ is expressed when both NFκB and IRF are activated or transfected than the sum of transcription with either factor alone (20, 21). Given the involvement of a chromatin structural protein HMG1 in bending the DNA, these findings led to a proposed model of an IFNβ “enhanceosome” (20), which recruits CBP (22) and requires the concerted activity of NFκB and IRF to recruit remodeling factors to move a nucleosome blocking the transcription start site (23). The IFNβ enhanceosome has been commonly understood to require two IRFs and NFκB to be bound to the enhancer for maximal transcriptional activation of IFNβ (20, 24, 25).

However, this model does not explain the regulation of IFNβ in all conditions. While IRF knockouts confirmed that IFNβ expression is dependent on IRF regardless of the PAMP stimulus or pathogen exposure (26–29), basal IRF may be sufficient in deregulated NFκB system conditions (30). Further, while dependence on NFκB was found upon stimulation with TLR4-inducing LPS (31, 32), NFκB is not required for IFNβ expression upon stimulation with TLR3- and RLR-inducing PolyIC or infection with Sendai virus (33, 27, 32, 34). Thus, it remains unclear what regulatory logic governs expression of this key innate immune-coordinating cytokine.

Here we developed a quantitative understanding of the regulatory logic by which IRF and NFκB govern IFNβ expression by evaluating alternate mathematical models of IFNβ regulatory enhancer states with published and newly generated datasets from a variety of stimulus and knockout conditions. We found that multiple synergy modes, each subject to particular dose-response relationships, are accessed in a condition-dependent manner to provide versatile but precise control of the IFNβ enhancer.

## RESULTS

### Defining a minimal combinatorial states model for IFNβ

To dissect the regulatory logic of the IFNβ enhancer, we used a combinatorial state ensemble model, which allows clear delineation of distinct regulatory modes (35, 36). To focus on the proposed NFκB-IRF synergy, we first modeled only two binding sites: one for NFκB and one for IRF (**Figure 1A, SI Methods**). The resulting two-site state ensemble model has four enhancer states: unbound, bound to IRF, bound to NFκB, or bound to both IRF&NFκB. The activities of the corresponding proteins in max-normalized units (MNU) for each state can be described by the functional state vector *S* and the transcriptional activity of each state by the vector *t*. The unbound state has no transcriptional activity (*t*_*0*_ *= 0*) and the IRF&NFκB state has maximal transcriptional activity (*t*_*IN*_ *= 1*), while the IRF state and the NFκB state have unknown transcriptional activities (*t*_*I*_ and *t*_*N*_, respectively). Each site has a binding equilibrium constant for its cognate protein, together comprising the vector *β*. As NFκB is an obligate dimer, we assume that it binds in one step with a binding equilibrium given by the parameter *K*_*N*_. In contrast, two IRF monomers may bind sequentially and thus show binding cooperativity (37, 38). To account for this, we let the binding equilibrium for IRF be 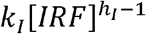, where *h* is a Hill coefficient that describes the potential cooperativity between IRF monomers. The promoter activity (*f*) is calculated from the *S, t*, and β vectors (**Figure 1B)**.

**Figure 1:**
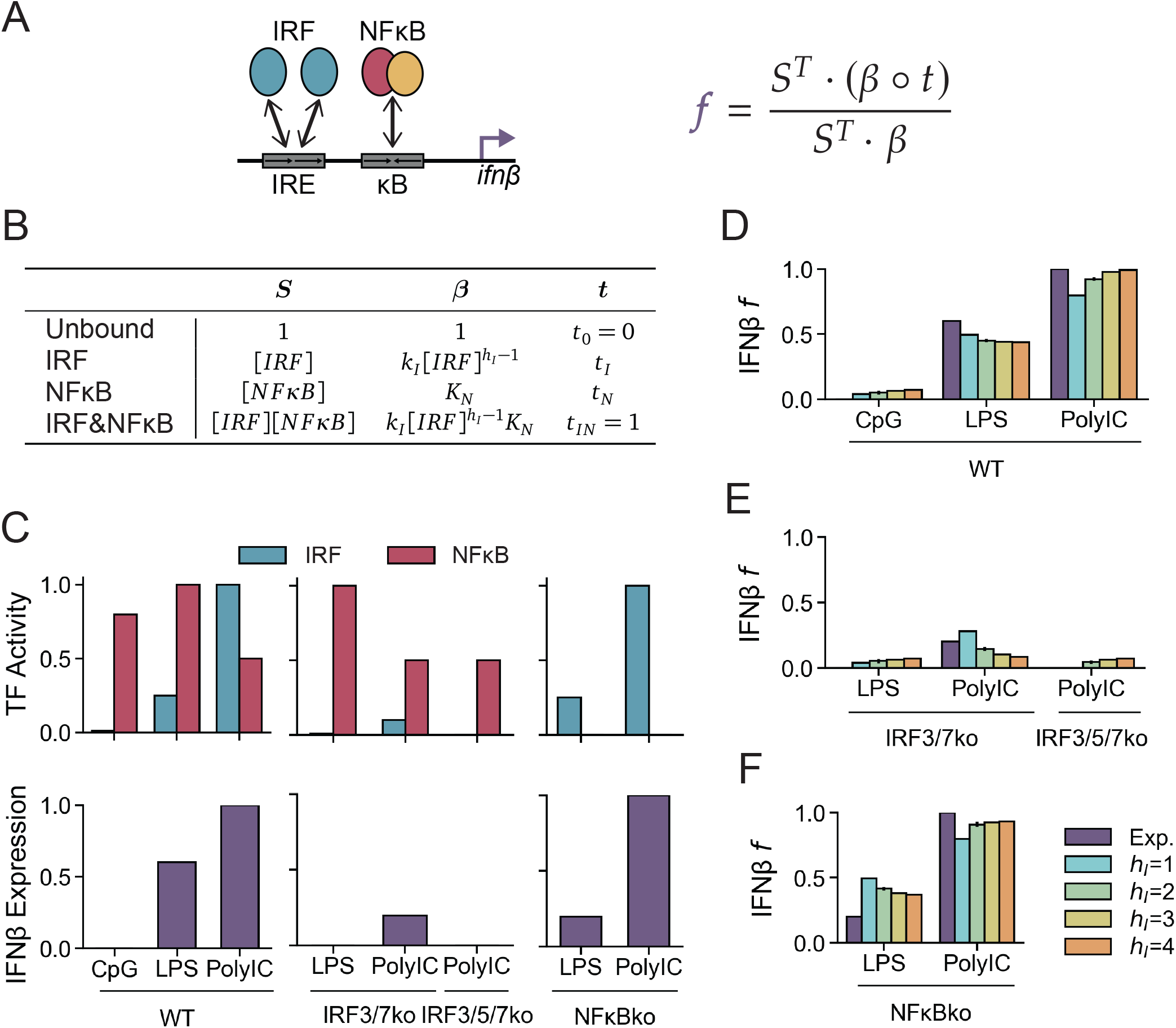
A two-site model does not account for the stimulus-specific NFκB requirement. (A) Schematic of IFNβ enhancer with two binding sites, one for each IRF and NFκB. Each protein has a corresponding binding affinity and activity. Function for predicting IFNβ mRNA is shown on top left. ∘ denotes elementwise multiplication. (B) Table of vectors for functional state (*S*), binding equilibrium constants (*β*), and transcriptional activity (*t*) for each state of the enhancer. (C) Experimental data showing max-normalized activities (MNU) of NFκB and IRF and max-normalized expression of IFNβ. (D-F) Experimental and simulated IFNβ expression for four models of *h*_*I*_, in the indicated conditions. Mean and standard deviation of the best 20 optimized parameter sets are shown for (D) WT, (E) IRF knockouts, (F) NFκB knockouts.

To fit the model, we surveyed literature data on IFNβ regulation to generate a data matrix (**Figure 1C, Table S1**) quantifying response to double-stranded CpG DNA, which activates NFκB but almost no IRF and stimulates no detectable IFNβ (16, 30), bacterial membrane component LPS, which induces low IRF and maximal NFκB and stimulates a medium amount of IFNβ (39, 40, 32, 31, 26), and viral analog PolyIC, which induces maximal IRF and stimulates maximal IFNβ (16, 41, 32, 27, 26, 29) in WT and four genetic knockouts. IRF3/5/7ko cells with PolyIC stimulus produce no detectable IFNβ, while IRF3/7ko cells with PolyIC stimulus produce a little IFNβ, though the weaker IRF-inducing stimulus LPS shows no IFNβ induction even in this double ko (26–29). In contrast, NFκBko cells, which lack subunits RelA and cRel, show severely diminished IFNβ expression when LPS is the stimulus, while they show no decrease in IFNβ expression from WT when PolyIC is the stimulus (33, 27, 34, 31, 32).

To determine the regulatory logic that best explains the experimental data, we fit the model parameters to the available data (**Figure 1C)** and selected the top 20 optimized parameter sets (**SI Methods**), which all gave essentially the same predicted values for IFNβ. For all *h*_*I*_ values, best-fit models correctly predicted IFNβ expression in WT cells (**Figure 1D**) and IRFkos (**Figure 1E**). However, all models failed to predict the deficiencies in IFNβ activation in NFκBko cells stimulated with LPS (**Figure 1F**). A high Hill coefficient allowed for more NFκB dependence in response to LPS than in response to PolyIC, but not to the extent observed experimentally and at the expense of LPS-inducibility in WT cells (**Figure 1D**). We were able to improve the fit to NFκB dependence by adding positive binding cooperativity (*C* parameter) between IRF and NFκB (**Figure S1A-B, Table S4**), but biochemical assays have shown that there is no such cooperativity (42). Additionally, we found that a NFκB Hill coefficient of 3 could also allow the model to fit the data (**Figure S1C-D, Table S3**)), but as NFκB is an obligate dimer and there is only one κB site in the IFNβ enhancer, this possibility is unlikely. Consequently, we concluded that the two-site state ensemble model is insufficient to explain the stimulus-specific NFκB dependence.

### A three-site model can account for the experimental data

We next tested a three-site model (**Figure 2A**), comprising a binding site for NFκB, a proximal IRF binding site (IRE_1_) and a distal IRF binding site (IRE_2_). IRE_1_ and IRE_2_ have binding equilibriums of 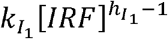 and 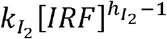 for their respective bound IRFs, where each *k*_*I*_ is a binding parameter and each *h*_*I*_ is a Hill coefficient reflecting binding cooperativity between IRF monomers. This model has eight states of binding: unbound, IRF bound to IRE_1_ (IRF_1_), IRF bound to IRE_2_ (IRF_2_), NFκB bound, IRF_1_&IRF_2,_ IRF_1_&NFκB, IRF_2_&NFκB, and IRF_1_&IRF_2_&NFκB. The transcriptional activities for the NFκB alone state (*t*_*N*_) and the IRF alone states (*t*_*I*_) are both unknown. New parameters are also defined for transcription from the states with two sites bound: 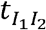 for the IRF_1_&IRF_2_ state, 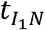 for the IRF_1_&NFκB state, and 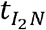 for the IRF_2_&NFκB state. These new parameters allow for functional synergy (e.g., 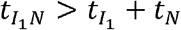) or antagonism between the different TFs. We identified optimized parameter sets for four different models, where IRF monomer cooperativity applies to neither IRF (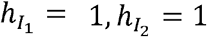; i.e., the 1&1 model), only the distal IRF (the 1&3 model), only the proximal IRF (the 3&1 model), or both (the 3&3 model).

**Figure 2:**
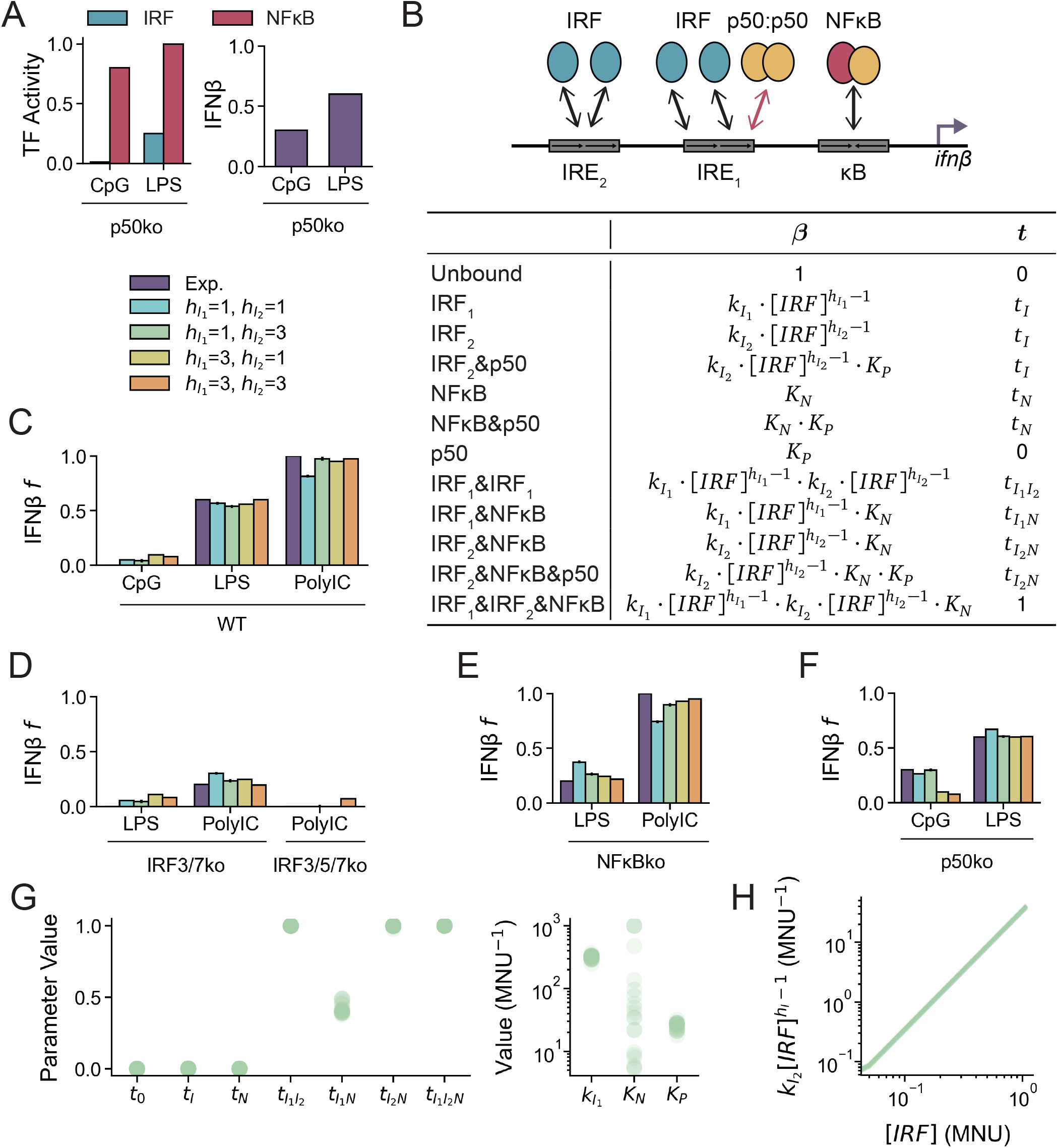
A three-site model accounts for the data in specific regulatory parameter regimes. (A) Left, schematic of three-site thermodynamic state model, with two IRE sites and one κB site. Each protein has a corresponding binding affinity and activity. Right, table of vectors for binding equilibrium constants (*β*) and transcriptional activity (*t*) for each state of the enhancer. (B-D) Experimental and simulated IFNβ expression for four models of 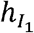 and 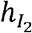, in the indicated conditions. Mean and standard deviation of the best 20 optimized parameter sets are shown for (B) WT, (C) IRF-dependent conditions, (D) NFκB-dependent conditions. (E) Ratio of experimental and simulated IFNβ expression in NFκBko to WT for LPS and PolyIC stimulus for all two-site and three-site models from best 20 optimized parameter sets.

All models adequately fit the data of stimulus-specific expression in wild type cells, though a high Hill coefficient for either IRF site improves the fit to PolyIC (**Figure 2B**). All models correctly predict deficiencies in IRFko cells, with a high Hill coefficient for both IRF sites (the 3&3 model) improving the fit (**Figure 2C**). Only the 1&3, 3&1, and 3&3 models closely recapitulate the NFκB dependence found in the literature (**Figure 2D**), showing a decrease in IFNβ in the NFκBko compared to WT under LPS stimulus not seen in the 1&1 model or any two-site model (**Figure 2E**). An NFκB Hill coefficient (*h*_*N*_) of 3, which is not supported by the known biochemistry, had minimal effect on the fits (**Figure S2**). We concluded that the three-site state ensemble model with sigmoidal binding of IRFs to IREs sufficiently accounts for the available IFNβ expression data.

### p50-homodimer tunes the IRF-NFκB regulatory logic

We next tested the three-site model with the previously reported finding that the p50:p50 homodimer binds the IRE_1_ binding site, competitively inhibiting IRF binding and IFNβ expression (30). In the absence of p50:p50, CpG, which does not activate IRF, is able to induce substantial IFNβ expression. However, the deficiency in p50:p50 results in only a minor increase in IFNβ with LPS (30). We added the p50ko condition to our collection of data (**Figure 3A**).

**Figure 3:**
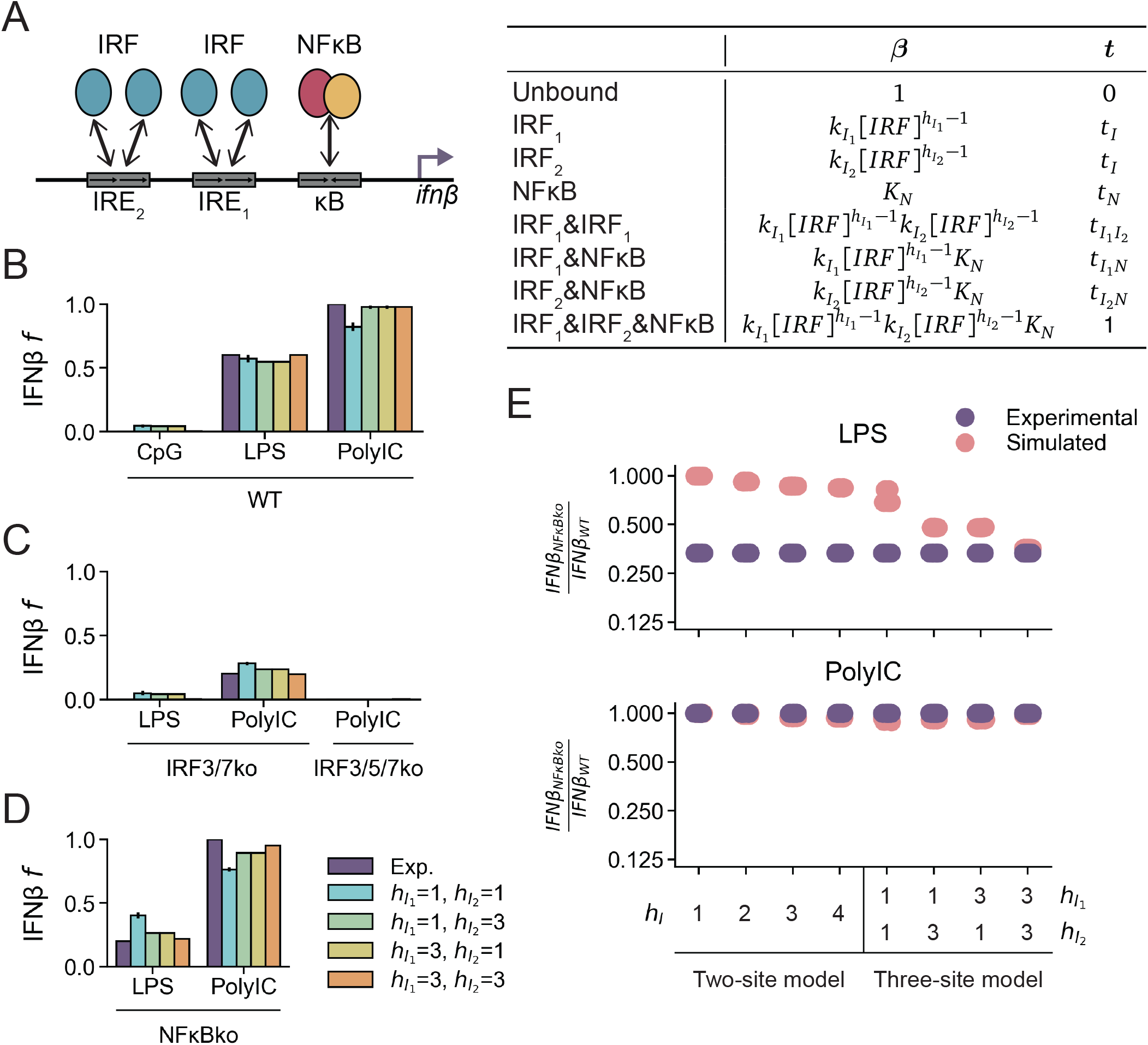
IRF responsiveness is tuned by the NFκB p50 homodimer. (A) Experimental data showing max-normalized activities (MNU) of NFκB and IRF and max-normalized expression of IFNβ in p50ko conditions. (B) Above, schematic of the three-site thermodynamic state model with competitive inhibition of IRF by p50:p50 homodimer at the proximal IRE site. Each protein has a corresponding binding affinity and activity. Below, table of vectors for binding equilibrium constants (*β*) and transcriptional activity (*t*) for each state of the enhancer. (C-F) Experimental and simulated IFNβ expression for four models of 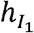 and 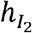 in the indicated conditions. Mean and standard deviation of the best 20 optimized parameter sets are shown for (C) WT, (D) IRF-dependent conditions, (E) NFκB-dependent conditions, (F) p50:p50-dependent conditions. (G) Best 20 optimized *t*, 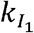, *K*_*N*_, and *K*_*P*_ parameter values for all data points in 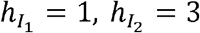 model. (H) Best 20 optimized IRF_2_ binding equilibrium as a function of IRF concentration in 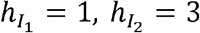 model.

We extended the three-site model to include p50:p50 vs. IRF competition for IRE_1_ (**Figure 3B**). p50:p50 may bind at the same time as IRF_2_ and/or NFκB, but not IRF_1_, resulting in four additional states of binding. p50:p50 does not contribute to transcription, such that transcription is determined only by any other bound TFs. We introduced the binding affinity of p50:p50 for IRE_1_ (*k*_*P*_) as a free parameter to be determined by the available experimental data. We fit parameters for four models (1&1, 1&3, 3&1, 3&3) and selected the top 20 optimized parameter sets. All models fit the WT data, though the 1&1 model had a noticeably worse fit to the WT PolyIC-stimulated condition (**Figure 3C**). IRF-dependent conditions were fit well across models, especially in the 1&3 and 3&1 models (**Figure 3D**). Again, the 1&1 model did not exhibit as much stimulus-specific NFκB dependence as the other models (**Figure 3E**). In the p50ko condition, while LPS expression was well-fit across all models, CpG expression was only fit by the 1&1 and 1&3 models (**Figure 3F**). Consequently, we proceeded with the 1&3 model, as it satisfactorily captured all observations.

The optimized parameters demonstrate high levels of functional synergy, where *t*_*I*_ and *t*_*N*_ are approximately 0 and 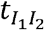 and 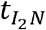 are approximately 1 (**Figure 3G, left**). There is somewhat less synergy between NFκB and the proximal IRF, as 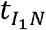 is around 0.5. Consequently, five of the twelve enhancer states (IRF_1_&IRF_2_, IRF_1_&NFκB, IRF_2_&NFκB, IRF_2_&NFκB&p50, and IRF_1_&IRF_2_&NFκB) are transcriptionally active, four of which (IRF_1_&IRF_2_, IRF_2_&NFκB, IRF_2_&NFκB&p50, and IRF_1_&IRF_2_&NFκB) are maximally active.

We next examined binding affinity in the 1&3 model. 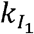 is an order of magnitude higher than *K*_*P*_ (≈300 MNU^-1^ compared to ≈25 MNU^-1^), which may be crucial for IRF to compete with p50:p50 when sufficient IRF is stimulated (**Figure 3G, right**). The maximal binding affinity for IRF_2_ 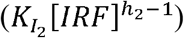, around 30 MNU^-1^, is likewise lower than 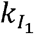 (**Figure 3H**). *K*_*N*_ is not well-determined by the data, as a range of values from 5 to 10^3^ MNU^-1^ satisfy the observations (**Figure 3G, right**). We concluded that the increased dynamic range of the IRF_2_ binding affinity is essential for capturing the high values of IFNβ in PolyIC-stimulated WT and NFκBko conditions. We conclude that a model where IRF_1_ has a binding affinity high enough to compete with p50:p50, IRF_2_ has sigmoidal binding, and all TF pairs have high synergy can explain all data.

### Alternate model conceptions show no improvement

We then investigated if binding cooperativity between IRF dimers would improve the fit of the model. We added a parameter, *C*, to the binding equilibrium of the IRF_1_&IRF_2_ state, which specified positive (*C* > 1) or negative binding cooperativity (*C* < 1) between IRF dimers at IRE_1_ and IRE_2_ (**Table S8, SI Methods**). While positive cooperativity slightly improved the fit of the 1&1 model, it did not affect the fit of the 1&3 model, which remained the only acceptable model (**Figures S3A-B**). We concluded that binding cooperativity is dispensable for fitting functional synergy. Next, we considered that functional synergy may plausibly be only possible between neighboring proteins and tested a model with 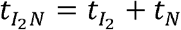 (**SI Methods**). We found that again only the 1&3 model was acceptable (**Figure S4A**). With distal synergy disallowed, 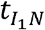 is approximately 1, so that all states yield either no transcription or maximal transcription (**Figure S4B**). As before, 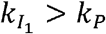 and 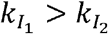 for all IRF activities (**Figure S4C**). Therefore, the only difference in this model is a higher 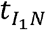, and neither alternate model improves the fit to the data.

### Modes of synergy explain stimulus specificities

Using the best-fitting 1&3 model, we investigated the probabilities of the 12 enhancer states under the 10 stimulus conditions (**Figure 4A-D**). In the inactive basal condition, the enhancer is primarily in the inactive p50 (54%) and NFκB&p50 (32%) states (**Figure 4A**), while upon CpG stimulation the NFκB&p50 state is dominant (81%) and probability of being in an active state remains low (10%). However, upon LPS stimulation, the probabilities of enhancer states are 25% for the highly active IRF_1_&IRF_2_&NFκB state, 46% for the partially active IRF_1_&NFκB state, and only 15% for the inactive NFκB&p50 state. In response to PolyIC, the enhancer is almost entirely in the IRF_1_&IRF_2_&NFκB state (82%). The differences in active state probabilities explain IFNβ stimulus-specificity.

**Figure 4:**
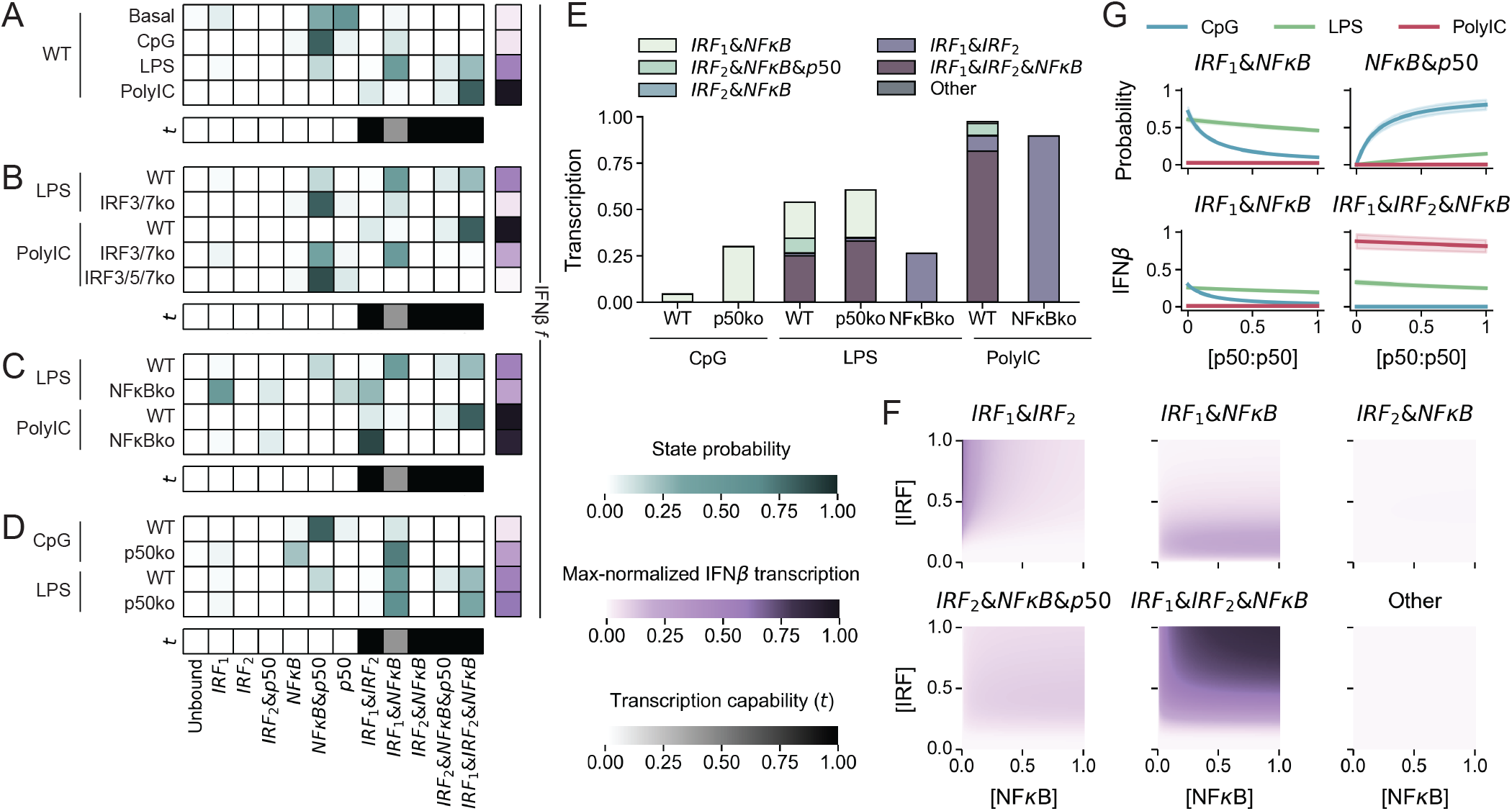
The enhancer state model reveals regulatory modes and can be used for forward predictions. (A-D) Heatmaps of the state probabilities (blue color bar) determined by the best-fit model. Bottom row (brown color bar) shows a heatmap of transcription capability (*t*) parameter for each state. Column on right (purple color bar) shows a heatmap of IFNβ *f* from model fit. State probability and *f* values are shown for: (A) WT in unstimulated and stimulated conditions, (B) IRF-deficient and corresponding WT conditions, (C) NFκB-deficient and corresponding WT conditions, (D) p50-deficient and corresponding WT conditions. (E) Bar plot of amount of transcription contributed by each state in stimulus-specific NFκB- and p50-dependent conditions determined by the best-fit model. (F) Heatmaps of the amount of transcription coming from each state for a range of NFκB and IRF input concentrations in WT predicted by the best-fit model. (G) Mean and standard deviation of predicted probability of the IRF_1_&NFκB and NFκB&p50 states (top) and IFNβ produced by the IRF_1_&NFκB and IRF_1_&IRF_2_&NFκB states (bottom) for different concentrations of p50:p50 under CpG, LPS, and PolyIC stimulus.

Reducing IRF activity perturbs the state distributions. In LPS stimulated IRF3/7ko cells, the probability of the IRF_1_&NFκB state decreases to 10%, leaving 82% for the NFκB&p50 state, explaining why almost all transcription is lost in this condition (**Figure 4B**). When the IRF3/7ko is stimulated with PolyIC, the partially active IRF_1_&NFκB state is accessed (48%), allowing the retention of some IFNβ production. This due to the high 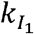 allowing even low concentrations of IRF to effectively compete with basally bound p50:p50. However, when all IRF activity is ablated in IRF3/5/7ko cells, the enhancer almost entirely occupies the inactive NFκB&p50 state (87%).

In LPS-stimulated NFκBko cells, the enhancer shifts not only from the IRF_1_&IRF_2_&NFκB state (25%) to the equally active IRF_1_&IRF_2_ state (26%) but also from the active IRF_1_&NFκB state (46%) into the inactive IRF_1_ state (49%), resulting in a decrease in IFNβ transcription (**Figure 4C**). Because PolyIC-stimulated WT cells specify primarily the IRF_1_&IRF_2_&NFκB state, NFκB-deficiency results in the equally active IRF_1_&IRF_2_ state (90%), explaining the NFκB independence in this condition.

While CpG does not activate IRF, in p50ko cells, basal IRF binding to the high affinity IRE_1_ and CpG-induced NFκB are sufficient for the enhancer to have a high probability (71%) of occupying the partially active IRF_1_&NFκB state (**Figure 4D**). However, in response to LPS, the probabilities of accessing the IRF_1_&NFκB or IRF_1_&IRF_2_&NFκB states only increase moderately over the WT condition (61% vs. 46% and 33% vs. 25%, respectively) given that LPS-induced IRF is sufficient to overcome the p50:p50 blockade in WT cells. This moderate increase in p50ko cells is associated with the loss of the IRF_2_&NFκB&p50 state.

To further understand the contribution of each state to IFNβ transcription in each condition, we examined the amount of max-normalized transcription units (TUs) contributed by each state (**Figure 4E**). In response to CpG, all transcription comes from the IRF_1_&NFκB state (0.04 TU), which increases in p50ko cells (0.30 TU). In response to LPS, transcription comes from IRF_1_&IRF_2_&NFκB (0.25 TU), IRF_1_&NFκB (0.19 TU), and IRF_1_&IRF_2_ (0.01 TU) states. Transcription from each of these states increases moderately when p50:p50 is absent (0.26, 0.33, and 0.02 TU, respectively), given that the IRF_2_&NFκB&p50 state is eliminated.

In NFκBko cells, the only available active state is IRF_1_&IRF_2_ (**Figure 4E**). In response to LPS, transcription from this state increases (from 0.01 to 0.26 TUs), reflecting the sum of the transcription from the IRF_1_&IRF_2_ and the IRF_1_&IRF_2_&NFκB states in WT. However, this does not compensate for the loss of the IRF_1_&NFκB state (from 0.19 TU to 0 TUs), causing an overall decrease in transcription. In response to PolyIC, most transcription is from the IRF_1_&IRF_2_ (0.08 TU) and the IRF_1_&IRF_2_&NFκB (0.81 TU) states, which converges on the IRF_1_&IRF_2_ state in the absence of NFκB (0.90 TU). All other active states show only a negligible loss of transcription in the NFκBko (0.08 TU). Overall, the probability distributions of active states explain the stimulus-specific and knockout condition-specific transcriptional control of IFNβ.

### Using the model for forward predictions

To use the combinatorial states model for forward predictions of biological scenarios not directly represented by the training data, we first asked whether the primary conclusions and best-fit parameters were robust to variations in the training data due to uncertainties in measurements. We generated 100 synthetic datasets with different levels of noise added to the NFκB and IRF input values and optimized parameters as before to the synthetic datasets. When noise corresponded to 1% error, best fit *t* parameters matched those in the original data, and best fit *k* parameters followed the same patterns as the original with a slightly wider range of values (**Figure S5A**). With 10% (**Figure S5B**) and 20% (**Figure S5C**) error, all best fit *t* parameters continued to match the original data except for 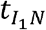, which had an increased upward spread towards 1. Only with 40% error did 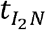 and 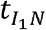 have high variation (**Figure S5D**). Importantly, we found that even at 20% noise, synergy was still required between the two IRFs as well as each IRF and NFκB (i.e., at least two sites must be bound to initiate transcription, **Figure S5C**). We conclude that 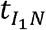 is the least robust parameter and that the synergy requirement of IRF_1_&NFκB may be less robust than that of IRF_1_&IRF_2_ or IRF_2_&NFκB, but that NFκB is always dispensable when IRF_1_ and IRF_2_ are bound. These studies indicated that our conclusions are robust to error in the assembled data matrix, providing confidence in the reliability of forward predictions in conditions outside of the training dataset.

We next undertook comprehensive dose response studies, predicting IFNβ transcription for a range of NFκB and IRF activities (**Figure 4F**). With enough IRF (>0.2 MNU) and non-zero NFκB, the IRF_1_&IRF_2_&NFκB state produces a large amount of IFNβ transcription and the IRF_2_&NFκB&p50 state contributes some IFNβ transcription. With enough IRF and low NFκB, the IRF_1_&IRF_2_ state produces moderate amounts of IFNβ transcription. With low but non-zero IRF, the IRF_1_&NFκB state produces most IFNβ transcription. All other states contribute very little to IFNβ transcription under any NFκB and IRF activities. We also predicted IFNβ transcription in p50ko cells (**Figure S4D**). The lower the concentration of IRF, the stronger the repressive effect of p50:p50. We further explored stimulus specificity as a function of p50:p50, finding that increased [p50:p50] decreased the probability of and transcription from the IRF_1_&NFκB state and increased the probability of the NFκB&P50 state under CpG and LPS stimulus while mildly decreasing transcription from the IRF_1_&IRF_2_&NFκB state under LPS and PolyIC stimulus (**Figure 4G**). These analyses reveal how IFNβ transcription can be tuned by different stimuli and cellular conditions that involve different IRF and NFκB activity levels.

## DISCUSSION

We report here the development of a quantitative model of the regulatory logic that governs IFNβ expression. It recapitulates a variety of stimulus-response data exploring the stimulus-specificity of IFNβ expression, its dependence on IRF and NFκB activators, and the role of the repressor p50:p50. We tested alternate models that are both qualitatively and quantitatively distinct in how the three transcription factors function together, synergistically or competitively, to control IFNβ expression. Even with limited data, we rejected many different model formulations, including various two-site and three-site models, and converged on specific regulatory insights. The resulting model reveals that each binding site and TF has a non-redundant regulatory role that is governed by quantitative specifications of its dose-response curve as well as functional interactions with partnering TFs on the enhancer. The specific role of each binding site explains why the enhancer sequence is so highly evolutionarily conserved and has no single nucleotide polymorphisms with a prevalence above 0.03% on dbSNP (43). Furthermore, the model had highly constrained parameters for IRF and p50 binding and for transcription capability of each state that were robust to potential measurement error.

We found a higher binding affinity for IRE_1_ than for IRE_2_ in all acceptable models. This is in line with biochemical studies, which have noted that the first half-site of IRE_1_ is GAGA, while IRE_2_ contains the suboptimal AAAA (42) and a 2 bp spacing between half-sites (44). The model also required a non-linear ultrasensitive dose response curve (i.e. a Hill coefficient above 1) at IRE_2_, but not at IRE_1_, where basal p50:p50 prevents spurious IRF binding. This agrees with biochemical studies that determined that IRF monomers bind cooperatively with each other and possibly with neighboring (constitutively present) AP1 at IRE_2_ (37). This remarkable agreement between the model inferred from stimulus-response data and biochemical findings not used for model fitting provides confidence in the model and offers support for the functional relevance of prior biochemical studies.

Given that all known IRF3-inducing stimuli also activate NFκB, NFκB is active in every WT pathogen-response condition. Each of these conditions yield NFκB bound to the enhancer, but whether NFκB binding is functionally important depends on context. Alone, it is unable to drive transcription of IFNβ, and when both IRF_1_ and IRF_2_ are present it is dispensable. Therefore, NFκB plays a functional role only when a single IRF dimer is present. Given that IRF has a higher binding affinity for IRE_2_ than for IRE_1_ and p50:p50 competes with IRF at IRE_2_, we may speculate that only in the case of very high p50:p50 homodimer levels might NFκB functionally synergize with the distal IRE_2_ to activate IFNβ expression. p50 homodimer concentration is context specific; due to the feedforward loop regulating NFκB dimers (45), cells pre-exposed to stimuli can contain higher levels of p50:p50 than naïve cells, reducing the contribution of the IRF_1_&NFκB state.

Previous research has not fully delineated the molecular mechanism that mediates transcription factor synergy. The simplest explanation, that IRF and NFκB have cooperativity in binding, is unsupported by the literature. In the atomic structure of the IFNβ enhancer, the two bound IRFs and NFκB do not have direct contacts (42, 37, 46). NFκB and the proximal IRF have no binding cooperativity when measured by EMSA, and the two IRF dimers may even be anti-cooperative (42). Instead, synergy may come from a subsequent mechanistic step, perhaps involving the recruitment of the cofactor CBP/p300 (47). CBP has disordered, flexible regions (48) and can adapt to different distances between the binding domains of IFNβ (42), supporting functional synergy between NFκB and either IRF. An alternate mechanism for synergy may be kinetic in which different regulatory steps of pre-initiation complex formation and initiation and elongation are determined by different TFs, but given that two IRFs are sufficient, such an underlying mechanism is unlikely.

Extending the model in the future can address different biological scenarios. Our model reflects pathogen sensing via PRR; future work could consider the regulation of IFNβ half-life (49), lower dependence on IRF3 and IRF7 (50), and roles of upstream enhancer elements (51) that occur during live infection. Data from additional biological conditions would not bring back models that were excluded by the present study, but it may add complexity to reveal intricacies of IFNβ regulation that are not yet captured. Similarly, the present model considers only IRF and NFκB as the key regulators of IFNβ expression, as early studies noted that AP1 and NFκB activation is always coordinated (52–55) and interdependent (56), and more recent studies described AP1’s function as a marker for open chromatin rather than a transcriptional activator (57, 58). However, additional data may necessitate the explicit consideration of AP1 in the regulatory logic of IFNβ expression.

The quantitative model of IFNβ regulation presented here contributes to 40 years of research into the regulation of the IFNβ enhancer. Though models of immune-response gene expression have advanced (59–64), IFNβ has long been a bottleneck in the development of interferon response models (65, 66). The present model for IFNβ promoter activity may therefore be a building block for the development of more comprehensive models that connect pathogen response signaling pathways with the large ISG expression program, expanding their utility for studies of IFNβ-related diseases as well, as molecular mechanisms for many inborn IFNβ deficiencies have not yet been identified (10).

## MATERIALS AND METHODS

Data collection is described in SI Appendix. Architecture of models and parameter sampling is described in the SI Appendix. Parameters were optimized using the Scipy v1.11.4 (67) implementation of the Nelder Mead algorithm.

## ACKNOWLEDGEMENTS

We thank the members of the Signaling Systems Lab for critical discussion and particularly thank Xiaolu Guo for valuable advice. We also thank Atef Ali for reading the manuscript. This work was funded by National Institutes of Health grants R01AI173214 and R01AI185026. This material is based upon work supported by the National Science Foundation Graduate Research Fellowship Program under Grant No. DGE-2034835. Any opinions, findings, and conclusions or recommendations expressed in this material are those of the authors and do not necessarily reflect the views of the National Science Foundation.

## REFERENCES

1. L. B. Ivashkiv, L. T. Donlin, Regulation of type I interferon responses. Nat. Rev. Immunol. 14, 36–49 (2014).

2. F. McNab, K. Mayer-Barber, A. Sher, A. Wack, A. O’Garra, Type I interferons in infectious disease. Nat. Rev. Immunol. 15, 87–103 (2015).

3. A. J. Sadler, B. R. G. Williams, Interferon-inducible antiviral effectors. Nat. Rev. Immunol. 8, 559–568 (2008).

4. C. L. Wilder, et al., A stimulus-contingent positive feedback loop enables IFN-β dose-dependent activation of pro-inflammatory genes. Mol. Syst. Biol. 19, e11294 (2023).

5. J. Crouse, U. Kalinke, A. Oxenius, Regulation of antiviral T cell responses by type I interferons. Nat. Rev. Immunol. 15, 231–242 (2015).

6. H. Jiang, et al., Interferon beta-1b reduces interferon gamma-induced antigen-presenting capacity of human glial and B cells. J. Neuroimmunol. 61, 17–25 (1995).

7. L. M. Snell, T. L. McGaha, D. G. Brooks, Type I Interferon in Chronic Virus Infection and Cancer. Trends Immunol. 38, 542–557 (2017).

8. W.-X. Sin, P. Li, J. P.-S. Yeong, K.-C. Chin, Activation and regulation of interferon-β in immune responses. Immunol. Res. 53, 25–40 (2012).

9. Y. J. Crow, D. B. Stetson, The type I interferonopathies: 10 years on. Nat. Rev. Immunol. 22, 471–483 (2022).

10. Y. J. Crow, J.-L. Casanova, Human life within a narrow range: The lethal ups and downs of type I interferons. Sci. Immunol. 9, eadm8185 (2024).

11. S. Makris, M. Paulsen, C. Johansson, Type I Interferons as Regulators of Lung Inflammation. Front. Immunol. 8 (2017).

12. G. Trinchieri, Type I interferon: friend or foe? J. Exp. Med. 207, 2053–2063 (2010).

13. K. Paucker, K. Cantell, W. Henle, Quantitative studies on viral interference in suspended L cells: III. Effect of interfering viruses and interferon on the growth rate of cells. Virology 17, 324–334 (1962).

14. H. Kumar, T. Kawai, S. Akira, Pathogen Recognition by the Innate Immune System. Int. Rev. Immunol. 30, 16–34 (2011).

15. L. Sun, J. Wu, F. Du, X. Chen, Z. J. Chen, Cyclic GMP-AMP Synthase Is a Cytosolic DNA Sensor That Activates the Type I Interferon Pathway. Science 339, 786–791 (2013).

16. S. Akira, K. Takeda, Toll-like receptor signalling. Nat. Rev. Immunol. 4, 499–511 (2004).

17. D. Goubau, S. Deddouche, C. Reis e Sousa, Cytosolic Sensing of Viruses. Immunity 38, 855–869 (2013).

18. D. Li, M. Wu, Pattern recognition receptors in health and diseases. Signal Transduct. Target. Ther. 6, 1–24 (2021).

19. K.-P. Hopfner, V. Hornung, Molecular mechanisms and cellular functions of cGAS–STING signalling. Nat. Rev. Mol. Cell Biol. 21, 501–521 (2020).

20. D. Thanos, T. Maniatis, Virus induction of human IFNβ gene expression requires the assembly of an enhanceosome. Cell 83, 1091–1100 (1995).

21. T. K. Kim, T. Maniatis, The Mechanism of Transcriptional Synergy of an In Vitro Assembled Interferon-β Enhanceosome. Mol. Cell 1, 119–129 (1997).

22. J. Yie, K. Senger, D. Thanos, Mechanism by which the IFN-beta enhanceosome activates transcription. Proc. Natl. Acad. Sci. U. S. A. 96, 13108–13113 (1999).

23. T. Agalioti, et al., Ordered Recruitment of Chromatin Modifying and General Transcription Factors to the IFN-β Promoter. Cell 103, 667–678 (2000).

24. D. Thanos, Mechanisms of transcriptional synergism of eukaryotic genes. The interferon-beta paradigm. Hypertens. Dallas Tex 1979 27, 1025–1029 (1996).

25. N. Munshi, et al., The IFN-β Enhancer: A Paradigm for Understanding Activation and Repression of Inducible Gene Expression. Cold Spring Harb. Symp. Quant. Biol. 64, 149–160 (1999).

26. S. Sakaguchi, et al., Essential role of IRF-3 in lipopolysaccharide-induced interferon-β gene expression and endotoxin shock. Biochem. Biophys. Res. Commun. 306, 860–866 (2003).

27. K. L. Peters, H. L. Smith, G. R. Stark, G. C. Sen, IRF-3-dependent, NFκB- and JNK-independent activation of the 561 and IFN-β genes in response to double-stranded RNA. Proc. Natl. Acad. Sci. 99, 6322–6327 (2002).

28. K. Honda, et al., IRF-7 is the master regulator of type-I interferon-dependent immune responses. Nature 434, 772–777 (2005).

29. H. M. Lazear, et al., IRF-3, IRF-5, and IRF-7 Coordinately Regulate the Type I IFN Response in Myeloid Dendritic Cells Downstream of MAVS Signaling. PLOS Pathog. 9, e1003118 (2013).

30. C. S. Cheng, et al., The Specificity of Innate Immune Responses Is Enforced by Repression of Interferon Response Elements by NF-κB p50. Sci. Signal. 4 (2011).

31. X. Wang, et al., Differential Requirement for the IKKβ/NF-κB Signaling Module in Regulating TLR-versus RLR-Induced Type 1 IFN Expression in Dendritic Cells. J. Immunol. 193, 2538–2545 (2014).

32. D. N. Rios, “Towards a Quantitative Understanding of NF-kappaB and Interferon Signaling During the Cellular Response to Pathogens /,” UC San Diego. (2014).

33. X. Wang, et al., Lack of Essential Role of NF-κB p50, RelA, and cRel Subunits in Virus-Induced Type 1 IFN Expression1. J. Immunol. 178, 6770–6776 (2007).

34. J. Wang, et al., NF-κB RelA Subunit Is Crucial for Early IFN-β Expression and Resistance to RNA Virus Replication. J. Immunol. 185, 1720–1729 (2010).

35. N. E. Buchler, U. Gerland, T. Hwa, On schemes of combinatorial transcription logic. Proc. Natl. Acad. Sci. 100, 5136–5141 (2003).

36. M. S. Sherman, B. A. Cohen, Thermodynamic State Ensemble Models of cis-Regulation. PLoS Comput. Biol. 8, e1002407 (2012).

37. D. Panne, T. Maniatis, S. C. Harrison, Crystal structure of ATF-2/c-Jun and IRF-3 bound to the interferon-beta enhancer. EMBO J. 23 (2004).

38. A. I. Dragan, V. V. Hargreaves, E. N. Makeyeva, P. L. Privalov, Mechanisms of activation of interferon regulator factor 3: the role of C-terminal domain phosphorylation in IRF-3 dimerization and DNA binding. Nucleic Acids Res. 35, 3525–3534 (2007).

39. Y.-C. Lu, W.-C. Yeh, P. S. Ohashi, LPS/TLR4 signal transduction pathway. Cytokine 42, 145–151 (2008).

40. K. Honda, A. Takaoka, T. Taniguchi, Type I Inteferon Gene Induction by the Interferon Regulatory Factor Family of Transcription Factors. Immunity 25, 349–360 (2006).

41. M. Matsumoto, T. Seya, TLR3: Interferon induction by double-stranded RNA including poly(I:C). Adv. Drug Deliv. Rev. 60, 805–812 (2008).

42. D. Panne, T. Maniatis, S. C. Harrison, An Atomic Model of the Interferon-β Enhanceosome. Cell 129, 1111–1123 (2007).

43. S. T. Sherry, et al., dbSNP: the NCBI database of genetic variation. Nucleic Acids Res. 29, 308–311 (2001).

44. C. R. Escalante, J. Yie, D. Thanos, A. K. Aggarwal, Structure of IRF-1 with bound DNA reveals determinants of interferon regulation. Nature 391, 103–106 (1998).

45. M. O. Metzig, et al., An incoherent feedforward loop interprets NFκB/RelA dynamics to determine TNFLJinduced necroptosis decisions. Mol. Syst. Biol. 16, e9677 (2020).

46. C. R. Escalante, E. Nistal-Villán, L. Shen, A. García-Sastre, A. K. Aggarwal, Structure of IRF-3 Bound to the PRDIII-I Regulatory Element of the Human Interferon-β Enhancer. Mol. Cell 26, 703–716 (2007).

47. M. Merika, A. J. Williams, G. Chen, T. Collins, D. Thanos, Recruitment of CBP/p300 by the IFN beta enhanceosome is required for synergistic activation of transcription. Mol. Cell 1, 277–287 (1998).

48. C. H. Lin, et al., A Small Domain of CBP/p300 Binds Diverse Proteins: Solution Structure and Functional Studies. Mol. Cell 8, 581–590 (2001).

49. M. M. Gaidt, et al., Self-guarding of MORC3 enables virulence factor-triggered immunity. Nature 600, 138–142 (2021).

50. S. Daffis, M. S. Suthar, K. J. Szretter, M. G. Jr, M. S. Diamond, Induction of IFN-β and the Innate Antiviral Response in Myeloid Cells Occurs through an IPS-1-Dependent Signal That Does Not Require IRF-3 and IRF-7. PLOS Pathog. 5, e1000607 (2009).

51. A. R. Banerjee, Y. J. Kim, T. H. Kim, A novel virus-inducible enhancer of the interferon-β gene with tightly linked promoter and enhancer activities. Nucleic Acids Res. 42, 12537–12554 (2014).

52. A. von Knethen, D. Callsen, B. Brüne, NF-κB and AP-1 Activation by Nitric Oxide Attenuated Apoptotic Cell Death in RAW 264.7 Macrophages. Mol. Biol. Cell 10, 361–372 (1999).

53. S.-W. Lee, S.-I. Han, H.-H. Kim, Z.-H. Lee, TAK1-dependent Activation of AP-1 and c-Jun N-terminal Kinase by Receptor Activator of NF-κB. BMB Rep. 35, 371–376 (2002).

54. C. Abate, L. Patel, F. J. Rauscher, T. Curran, Redox Regulation of Fos and Jun DNA-Binding Activity in Vitro. Science 249, 1157–1161 (1990).

55. M. B. Toledano, W. J. Leonard, Modulation of transcription factor NF-kappa B binding activity by oxidation-reduction in vitro. Proc. Natl. Acad. Sci. U. S. A. 88, 4328–4332 (1991).

56. S. Fujioka, et al., NF-κB and AP-1 Connection: Mechanism of NF-κB-Dependent Regulation of AP-1 Activity. Mol. Cell. Biol. 24, 7806–7819 (2004).

57. R. I. Martínez-Zamudio, et al., AP-1 imprints a reversible transcriptional programme of senescent cells. Nat. Cell Biol. 22, 842–855 (2020).

58. T. Vierbuchen, et al., AP-1 Transcription Factors and the BAF Complex Mediate Signal-Dependent Enhancer Selection. Mol. Cell 68, 1067-1082.e12 (2017).

59. M. Gilchrist, et al., Systems biology approaches identify ATF3 as a negative regulator of Toll-like receptor 4. Nature 441, 173–178 (2006).

60. M. Jovanovic, et al., Immunogenetics. Dynamic profiling of the protein life cycle in response to pathogens. Science 347, 1259038 (2015).

61. V. L. Bass, V. C. Wong, M. E. Bullock, S. Gaudet, K. Miller-Jensen, TNF stimulation primarily modulates transcriptional burst size of NF-κB-regulated genes. Mol. Syst. Biol. 17, e10127 (2021).

62. R. E. C. Lee, S. R. Walker, K. Savery, D. A. Frank, S. Gaudet, Fold change of nuclear NF-κB determines TNF-induced transcription in single cells. Mol. Cell 53, 867–879 (2014).

63. C. S. Cheng, et al., Iterative Modeling Reveals Evidence of Sequential Transcriptional Control Mechanisms. Cell Syst. 4, 330-343.e5 (2017).

64. S. Sen, Z. Cheng, K. M. Sheu, Y. H. Chen, A. Hoffmann, Gene Regulatory Strategies that Decode the Duration of NFκB Dynamics Contribute to LPS-versus TNF-Specific Gene Expression. Cell Syst. 10, 169-182.e5 (2020).

65. F. Kok, et al., Disentangling molecular mechanisms regulating sensitization of interferon alpha signal transduction. Mol. Syst. Biol. 16, e8955 (2020).

66. Z. Korwek, et al., Nonself RNA rewires IFN-β signaling: A mathematical model of the innate immune response. Sci. Signal. 16, eabq1173 (2023).

67. P. Virtanen, et al., SciPy 1.0: Fundamental Algorithms for Scientific Computing in Python. Nat. Methods 17, 261–272 (2020).

