## Supplementary Materials for "Gene regulatory logic of the interferon-β enhancer contains multiple selectively deployed modes of transcription factor synergy"

### The cis-regulatory logic of the type I interferon enhancer allows for tunable, stimulus-specific responses

**This PDF file includes:**

Supporting text
Figs. S1 to S5
Tables S1 to S8
SI References

### Supporting Information Text

### Methods

**Data Collection.** We assembled our data from a review of literature and some of our own measurements. From past literature on receptor mechanisms (1–4), we established the following:

- CpG activates NFκB
- LPS activates NFκB and some IRF
- PolyIC activates IRF and some NFκB

From our understanding of the IRF and NFκB families as well as data on IRF and NFκB knockouts (5), we determined the following:

- The IRF3/7ko reduces the amount of IRF activated by LPS to negligible levels. The amount of NFκB activated by LPS is slightly increased due to compensation.
- The IRF3/7ko reduces the amount of IRF activated by PolyIC, but some IRF5 is still activated. The IRF3/7ko completely removes any activated IRF.
- The RelA/cRelko (NFκBko) eliminates activated NFκB without affecting IRF under LPS and PolyIC stimuli.

There is substantial literature measuring the amount of IFNβ produced from various stimuli and genotypes in murine fibroblasts, dendritic cells, and macrophages. We observed the following:

- The IRF3/7ko eliminates IFNβ expression with LPS stimulus and reduces IFNβ expression with PolyIC (5, 6), and other IRFkos and stimuli show similar trend (7–10).
- The NFκBko reduces IFNβ expression with LPS stimulus while minimally affecting IFNβ expression with PolyIC stimulus (5, 11–14).

Finally, our lab has found that the p50ko substantially increases IFNβ with CpG stimulus, but not with LPS stimulus (15).
We quantitated all these these observations into Table S1.

**Table S1. Data collected and used for fitting. Data is shown in Figure 1.**

| Genotype | Stimulus | IRF | NFkB | IFNb | p50 |
|---|---|---|---|---|---|
| WT | LPS | 0.25 | 1 | 0.6 | 1 |
| p50KO | LPS | 0.25 | 1 | 0.6 | 0 |
| WT | CpG | 0.01 | 0.8 | 0 | 1 |
| p50KO | CpG | 0.01 | 0.8 | 0.3 | 0 |
| relacrelKO | LPS | 0.25 | 0 | 0.2 | 1 |
| irf3irf7KO | LPS | 0.01 | 1 | 0 | 1 |
| WT | polyIC | 1 | 0.5 | 1 | 1 |
| relacrelKO | polyIC | 1 | 0 | 1 | 1 |
| irf3irf7KO | polyIC | 0.1 | 0.5 | 0.2 | 1 |
| irf3irf5irf7KO | polyIC | 0 | 0.5 | 0 | 1 |

### Two-Site Model

**Defining the model.** In the thermodynamic state model, the enhancer is modeled as a set of transcriptionally active states, where the probability of each state occurring is weighted by the amount of transcription promoted by that state ((16), reviewed in (17)). We defined a two-site model with a binding site for IRF and a binding site for NFκB. This enhancer can have four states: unbound, IRF, NFκB, and IRF&NFκB. Each state has a corresponding functional state, binding affinity, and transcriptional capability, represented by $S$, $\beta$, and $t$ respectively (Table S2).

**Table S2. Two-site model**

| | $\boldsymbol{S}$ | $\boldsymbol{\beta}$ | $\boldsymbol{t}$ |
|---|---|---|---|
| Unbound | $1$ | $1$ | $t_0$ |
| IRF | $[IRF]$ | $k_I[IRF]^{h_I-1}$ | $t_I$ |
| NFκB | $[NF\kappa B]$ | $k_N$ | $t_N$ |
| IRF&NFκB | $[IRF][NF\kappa B]$ | $k_I[IRF]^{h_I-1}k_N$ | $t_{IN}$ |

As IRF and NFκB are activators of transcription, the unbound state is assumed to promote no transcription ($t_0 = 0$) and the fully bound IRF&NFκB state is assumed to promote maximal transcription ($t_{IN} = 1$). [IRF] and [NFκB] are defined by the max-normalized activites of IRF and NFκB for each condition. $k_I$, $k_N$, $h_I$, $t_I$, and $t_N$ are free parameters.

From thermodynamic equations for proteins binding, we can write the function for the production of IFNβ ($f$) in terms of $S$, $\beta$, and $t$ (17) (Equation 1).

$$f = \frac{S^T \cdot (\beta \circ t)}{S^T \cdot \beta} \tag{1}$$

where $\circ$ represents element-wise multiplication and $\cdot$ represents the dot-product.

It can be useful to think of this equation in the form given in Equation 2. Here, $P$ is the vector containing the probability that the enhancer will occupy each state.

$$f = P \cdot t \tag{2}$$

$$P = \frac{S^T \circ \beta^T}{S^T \cdot \beta} \tag{3}$$

We can then calculate the probabilities of each of the states using Equation 3, yielding the following.

$$P(\text{Unbound}) = \frac{1}{1 + k_I[IRF]^{h_I} + k_N[NF\kappa B] + k_I[IRF]^{h_I}k_N[NF\kappa B]}$$

$$P(IRF) = \frac{k_I[IRF]^{h_I}}{1 + k_I[IRF]^{h_I} + k_N[NF\kappa B] + k_I[IRF]^{h_I}k_N[NF\kappa B]}$$

$$P(NF\kappa B) = \frac{k_N[NF\kappa B]}{1 + k_I[IRF]^{h_I} + k_N[NF\kappa B] + k_I[IRF]^{h_I}k_N[NF\kappa B]}$$

$$P(IRF\&NF\kappa B) = \frac{k_I[IRF]^{h_I}k_N[NF\kappa B]}{1 + k_I[IRF]^{h_I} + k_N[NF\kappa B] + k_I[IRF]^{h_I}k_N[NF\kappa B]}$$

**Fitting the model.** We fit the five parameters to the ten data points shown in Figure 1C-F. First, we generated a grid of all combinations of 11 evenly spaced parameters in the ranges $[0, 1]$ for $t_I$ and $t_N$ (on a linear scale) and in the range $[10^{-3}, 10^3]$ for $k_I$ and $k_N$ (on a logarithmic scale), giving $11^4$ total parameter sets. For each value of $h_I$ (1,2,3,4), we calculated IFNβ for the 11 data points using Equation 1 given each set of parameters in the grid. With the predicted IFNβ, we calculated the RMSD to the data IFNβ (from Figure 1D-F) and selected the 100 parameter sets with the lowest RMSD to use as initial values for optimization.

For optimization, we took each of the 100 initial parameter sets and minimized a loss function for RMSD (Equation 4) using the Scipy (v1.11.4) (18) implementation of the Nelder Mead algorithm.

$$\min_f \sqrt{\frac{1}{10} \sum_{c \in \text{conditions}} (f_c(\text{model}) - f_c(\text{exp}))^2} \tag{4}$$

This resulted in 100 optimized parameter sets for each $h_I$ value, many of which were virtually identical (difference on the order of magnitude of $< 10^{-5}$ or lower). We selected the 20 sets with the lowest RMSD.

**Supplementary models.** We tested the effect of a Hill coefficient on NFκB in the two-site model (Table S3). This model was fit as described above for each value of $h_N$ in 1,2,3,4.

**Table S3. Two-site model with NFκB cooperativity**

| | $S$ | $\beta$ | $t$ |
|---|---|---|---|
| Unbound | 1 | 1 | $t_0$ |
| IRF | $[IRF]$ | $k_I[IRF]^{h_I-1}$ | $t_I$ |
| NFκB | $[NF\kappa B]$ | $k_N[NF\kappa B]^{h_N-1}$ | $t_N$ |
| IRF&NFκB | $[IRF][NF\kappa B]$ | $k_I[IRF]^{h_I-1}k_N[NF\kappa B]^{h_N-1}$ | $t_{IN}$ |

We also modeled the enhancer with binding cooperativity, showing in Table S4. This model was fit as described above, with the addition of 11 evenly spaces values of $C$ were sampled from the range $[10^{-3}, 10^3]$ giving $11^5$ total parameter sets for the initial scan.

**Table S4. Two-site model with binding cooperativity**

| | $\boldsymbol{S}$ | $\boldsymbol{\beta}$ | $\boldsymbol{t}$ |
|---|---|---|---|
| Unbound | $1$ | $1$ | $t_0$ |
| IRF | $[IRF]$ | $k_I[IRF]^{h_I-1}$ | $t_I$ |
| NFκB | $[NF\kappa B]$ | $k_N$ | $t_N$ |
| IRF&NFκB | $[IRF][NF\kappa B]$ | $k_I[IRF]^{h_I-1}k_N C$ | $t_{IN}$ |

### Three-Site Model

**Defining the model.** In the three-site model, the three binding sites (NFκB, IRF, and IRF) are considered separately. There are 8 states, each with a corresponding component of the $S$, $\beta$, and $t$ vectors (Table S5). The model was constructed as with the two-site model. We let double-bound states have undetermined transcriptional capability when bound together, allowing for transcriptional synergy.

**Table S5. Three-site model**

| | $\boldsymbol{S}$ | $\boldsymbol{\beta}$ | $\boldsymbol{t}$ |
|---|---|---|---|
| None | $1$ | $1$ | $0$ |
| $IRF_1$ | $[IRF]$ | $k_{I_1}[IRF]^{h_{I_1}-1}$ | $t_I$ |
| $IRF_2$ | $[IRF]$ | $k_{I_2}[IRF]^{h_{I_2}-1}$ | $t_I$ |
| NFκB | $[NF\kappa B]$ | $k_N$ | $t_N$ |
| $IRF_1$&$IRF_2$ | $[IRF][IRF]$ | $k_{I_1}[IRF]^{h_{I_1}-1}k_{I_2}[IRF]^{h_{I_2}-1}$ | $t_{I_1 I_2}$ |
| $IRF_1$&NFκB | $[IRF][NF\kappa B]$ | $k_{I_1}[IRF]^{h_{I_1}-1}k_N$ | $t_{I_1 N}$ |
| $IRF_2$ &NFκB | $[IRF][NF\kappa B]$ | $k_{I_2}[IRF]^{h_{I_2}-1}k_N$ | $t_{I_2 N}$ |
| $IRF_1$&$IRF_2$&NFκB | $[IRF][IRF][NF\kappa B]$ | $k_{I_1}[IRF]^{h_{I_1}-1}k_{I_2}[IRF]^{h_{I_2}-1}k_N$ | $1$ |

**Fitting the model.** To fit the parameters to the data, we first generated a grid of $10^6$ pseudo-randomly sampled parameter values for $t_I$, $t_N$, $t_{I_1 I_2}$, $t_{I_1 N}$, and $t_{I_2 N}$ from a uniform distribution between 0 and 1 and $k_{I_1}$, $k_{I_2}$, and $k_N$ from a logarithmic distribution between $10^{-3}$ and $10^3$ using Latin Hypercube Sampling (LHS). LHS defines a grid with $10^6$ intervals for each parameter and randomly samples one point from each interval. This ensures an even distribution of sampled parameter values. For each combination of $h_{I_1}$ and $h_{I_2}$ (1&1, 1&3, 3&1, 3&3), we calculated IFNβ for the 11 data points using Equation 1 for each set of parameters. With the predicted IFNβ, we calculated the RMSD to the data IFNβ and selected the 100 parameter sets with the lowest RMSD. For optimization, we took each of the 100 initial parameter sets and minimized RMSD (Equation 4) using the Nelder Mead algorithm. This resulted in 100 optimized parameter sets for each combination of Hill values. We selected the 20 sets with the lowest RMSD.

### Three-Site Model with p50:p50 competition

**Defining the model.** When p50:p50 competition is added to the three-site model, p50:p50 is able to bind to the $IRF_1$ binding site in place of IRF. We assumed that the binding of p50 neither activated nor repressed transcription, so the transcriptional capability value is determined by all other bound proteins. The binding affinity of p50 ($k_p$) is unknown. The full model has 12 possible states (Table S6).

**Table S6. p50 model**

| | $\boldsymbol{S}$ | $\boldsymbol{\beta}$ | $\boldsymbol{t}$ |
|---|---|---|---|
| None | $1$ | $1$ | $0$ |
| $IRF_1$ | $[IRF]$ | $k_{I_1}[IRF]^{h_{I_1}-1}$ | $t_I$ |
| $IRF_2$ | $[IRF]$ | $k_{I_2}[IRF]^{h_{I_2}-1}$ | $t_I$ |
| $IRF_2$&p50 | $[IRF][p50]$ | $k_{I_2}[IRF]^{h_{I_2}-1}k_P$ | $t_I$ |
| NFκB | $[NF\kappa B]$ | $k_N$ | $t_N$ |
| NFκB&p50 | $[NF\kappa B][p50]$ | $k_N k_P$ | $t_N$ |
| p50 | $[p50]$ | $k_P$ | $0$ |
| $IRF_1$&$IRF_2$ | $[IRF][IRF]$ | $k_{I_1}[IRF]^{h_{I_1}-1}k_{I_2}[IRF]^{h_{I_2}-1}$ | $t_{I_1 I_2}$ |
| $IRF_1$&NFκB | $[IRF][NF\kappa B]$ | $k_{I_1}[IRF]^{h_{I_1}-1}k_N$ | $t_{I_1 N}$ |
| $IRF_2$ &NFκB | $[IRF][NF\kappa B]$ | $k_{I_2}[IRF]^{h_{I_2}-1}k_N$ | $t_{I_2 N}$ |
| $IRF_2$ &NFκB&p50 | $[IRF][NF\kappa B][p50]$ | $k_{I_2}[IRF]^{h_{I_2}-1}k_N k_P$ | $t_{I_2 N}$ |
| $IRF_1$&$IRF_2$&NFκB | $[IRF][IRF][NF\kappa B]$ | $k_{I_1}[IRF]^{h_{I_1}-1}k_{I_2}[IRF]^{h_{I_2}-1}k_N$ | $1$ |

**Fitting the model.** To fit the parameters to the data, we first generated a grid of $10^6$ pseudo-randomly sampled parameter values for $t_I$, $t_N$, $t_{I_1 I_2}$, $t_{I_1 N}$, and $t_{I_2 N}$ from a uniform distribution between 0 and 1 and $k_{I_1}$, $k_{I_2}$, $k_N$, and $k_P$ from a logarithmic distribution between $10^{-3}$ and $10^3$ using LHS. For each combination of $h_{I_1}$ and $h_{I_2}$ (1&1, 1&3, 3&1, 3&3), we calculated IFNβ for the 11 data points using Equation 1 for each set of parameters. With the predicted IFNβ, we calculated the RMSD to the data IFNβ and selected the 100 parameter sets with the lowest RMSD. For optimization, we took each of the 100 initial parameter sets and minimized RMSD (Equation 4) using the Nelder Mead algorithm. This resulted in 100 optimized parameter sets for each combination of Hill values. We selected the 20 sets with the lowest RMSD.

**Calculations.** The state probabilities were calculated for each of the top 20 parameter sets using Equation 3 and the mean was taken across these 20 sets. The transcription ($f_s$) for each state $s$ was calculated using Equation 5.

$$f_s = \frac{S_s \beta_s t_s}{S^T \cdot \beta} \qquad [5]$$

Variance of state probabilities and state transcription values among the 20 different optimized parameter sets was minimal.

**Supplementary models.** We tested the three-site model with p50 competition while enforcing a lack of synergy between NFκB and $\text{IRF}_2$, given by Table S7. Parameters were fit as described above.

**Table S7. p50 model without synergy between NFκB and $\text{IRF}_2$**

| | $\boldsymbol{S}$ | $\boldsymbol{\beta}$ | $\boldsymbol{t}$ |
|---|---|---|---|
| None | 1 | 1 | 0 |
| $\text{IRF}_1$ | $[IRF]$ | $k_{I_1}[IRF]^{h_{I_1}-1}$ | $t_I$ |
| $\text{IRF}_2$ | $[IRF]$ | $k_{I_2}[IRF]^{h_{I_2}-1}$ | $t_I$ |
| $\text{IRF}_2$&p50 | $[IRF][p50]$ | $k_{I_2}[IRF]^{h_{I_2}-1}k_P$ | $t_I$ |
| NFκB | $[NF\kappa B]$ | $k_N$ | $t_N$ |
| NFκB&p50 | $[NF\kappa B][p50]$ | $k_N k_P$ | $t_N$ |
| p50 | $[p50]$ | $k_P$ | 0 |
| $\text{IRF}_1$&$\text{IRF}_2$ | $[IRF][IRF]$ | $k_{I_1}[IRF]^{h_{I_1}-1}k_{I_2}[IRF]^{h_{I_2}-1}$ | $t_{I_1 I_2}$ |
| $\text{IRF}_1$&NFκB | $[IRF][NF\kappa B]$ | $k_{I_1}[IRF]^{h_{I_1}-1}k_N$ | $t_{I_1 N}$ |
| $\text{IRF}_2$ &NFκB | $[IRF][NF\kappa B]$ | $k_{I_2}[IRF]^{h_{I_2}-1}k_N$ | $t_{I_2}+t_N$ |
| $\text{IRF}_2$ &NFκB&p50 | $[IRF][NF\kappa B][p50]$ | $k_{I_2}[IRF]^{h_{I_2}-1}k_N k_P$ | $t_{I_2}+t_N$ |
| $\text{IRF}_1$&$\text{IRF}_2$&NFκB | $[IRF][IRF][NF\kappa B]$ | $k_{I_1}[IRF]^{h_{I_1}-1}k_{I_2}[IRF]^{h_{I_2}-1}k_N$ | 1 |

We also tested the three-site model with p50 competition with the addition of IRF binding cooperativity. In this model (Table S8), the binding affinity for states with both IRF binding sites bound by IRF had an additional $C$ parameter. The model was fit in the same manner as the three-site models, with $C$ being initially sampled between $10^{-3}$ and $10^3$.

**Table S8. p50 model with IRF binding cooperativity**

| | $\boldsymbol{S}$ | $\boldsymbol{\beta}$ | $\boldsymbol{t}$ |
|---|---|---|---|
| None | 1 | 1 | 0 |
| $\text{IRF}_1$ | $[IRF]$ | $k_{I_1}[IRF]^{h_{I_1}-1}$ | $t_I$ |
| $\text{IRF}_2$ | $[IRF]$ | $k_{I_2}[IRF]^{h_{I_2}-1}$ | $t_I$ |
| $\text{IRF}_2$&p50 | $[IRF][p50]$ | $k_{I_2}[IRF]^{h_{I_2}-1}k_P$ | $t_I$ |
| NFκB | $[NF\kappa B]$ | $k_N$ | $t_N$ |
| NFκB&p50 | $[NF\kappa B][p50]$ | $k_N k_P$ | $t_N$ |
| p50 | $[p50]$ | $k_P$ | 0 |
| $\text{IRF}_1$&$\text{IRF}_2$ | $[IRF][IRF]$ | $k_{I_1}[IRF]^{h_{I_1}-1}k_{I_2}[IRF]^{h_{I_2}-1}C$ | $t_{I_1 I_2}$ |
| $\text{IRF}_1$&NFκB | $[IRF][NF\kappa B]$ | $k_{I_1}[IRF]^{h_{I_1}-1}k_N$ | $t_{I_1 N}$ |
| $\text{IRF}_2$ &NFκB | $[IRF][NF\kappa B]$ | $k_{I_2}[IRF]^{h_{I_2}-1}k_N$ | $t_{I_2 N}$ |
| $\text{IRF}_2$ &NFκB&p50 | $[IRF][NF\kappa B][p50]$ | $k_{I_2}[IRF]^{h_{I_2}-1}k_N k_P$ | $t_{I_2 N}$ |
| $\text{IRF}_1$&$\text{IRF}_2$&NFκB | $[IRF][IRF][NF\kappa B]$ | $k_{I_1}[IRF]^{h_{I_1}-1}k_{I_2}[IRF]^{h_{I_2}-1}k_N C$ | 1 |

### Model analysis

**Robustness analysis.** For each error $e$ in 1%, 10%, 20%, and 40%, we generated 100 synthetic data sets containing sampled concentrations of NFκB and IRF for each of the 10 data points. For each data point, a value for [NFκB] and [IRF] was sampled from a normal distribution with mean $\mu = [NF\kappa B]_{\text{data}}$ and $\mu = [IRF]_{\text{data}}$, respectively, and standard deviation $\sigma = e$. Data points were rejected if they failed one of two constraints:

1. $[IRF]_{\text{LPS,WT}} > [IRF]_{\text{LPS,IRF3/7ko}}$
2. $[IRF]_{\text{PolyIC,IRF3/7ko}} > [IRF]_{\text{PolyIC,IRF3/5/7ko}}$

IFNβ was set to the same value as in the original data. For each synthetic dataset, parameters were fit as described above, and mean value of top 20 parameter fits was selected for each dataset.

**Other analysis.** State probabilities were calculated using Equation 3. Transcription given by each state was calculated from Equation 6, which gives a vector that can be summed to give the value of $f$.

$$f = \frac{S^T \circ (\beta \circ t)}{S^T \cdot \beta} \qquad [6]$$

To make forward predictions, we generated a grid of 50 of evenly spaced NFκB and IRF concentrations (2500 combinations) between 1 and 0. We used Equation 6 with WT p50:p50 (i.e., [p50]=1 in Table S6) to calculate the amount of transcription coming from each state at each NFκB and IRF combination for each of the top 20 parameter sets, then took the mean of state transcription across the 20 sets. This process was repeated with a p50ko condition ([p50]=0 in Table S6) .

To predict the effect of varying concentrations of p50:p50, we generated 101 evenly spaced [p50] values. We assumed that each stimulus activated the same amount of IRF and NFκB regardless of [p50], so we used [IRF] and [NFκB] values from Table S1 for CpG, LPS, and PolyIC stimulus with each [p50] value to calculate state probabilities and transcription. We made this calculation for each of the top 20 parameter sets, then took the mean and standard deviation of state transcription across the 20 sets.

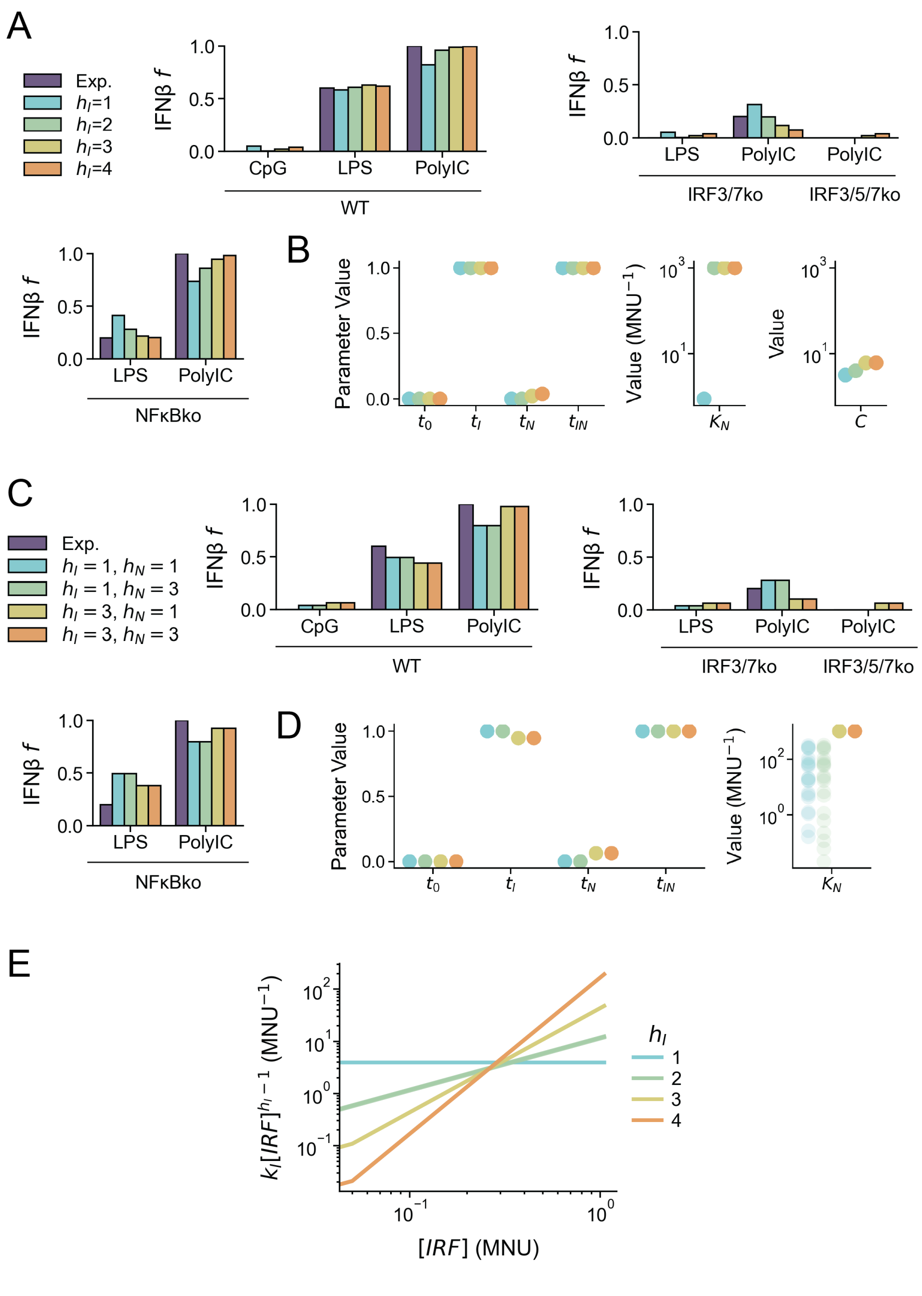


**Fig. S1.** (A) Experimental and mean predicted expression of IFNβ from best 20 optimized parameter sets for all data points using four models of $h_I$ for two-site model with binding cooperativity. (B) Best 20 optimized $t$, $K_N$, and $C$ parameter values for all data points for two-site model with binding cooperativity represented by scaling factor $C$. (C) Experimental and mean predicted expression of IFNβ from best 20 optimized parameter sets for all data points using four models of $h_I$ and $h_N$ in two-site model. (D) Best 20 optimized $t$ and $K_N$ parameter values for all data points for four models of $h_I$ and $h_N$ in two-site model. (E) Value of IRF binding equilibrium as a function of $[IRF]$ for four models of $h_I$ in two-site model.

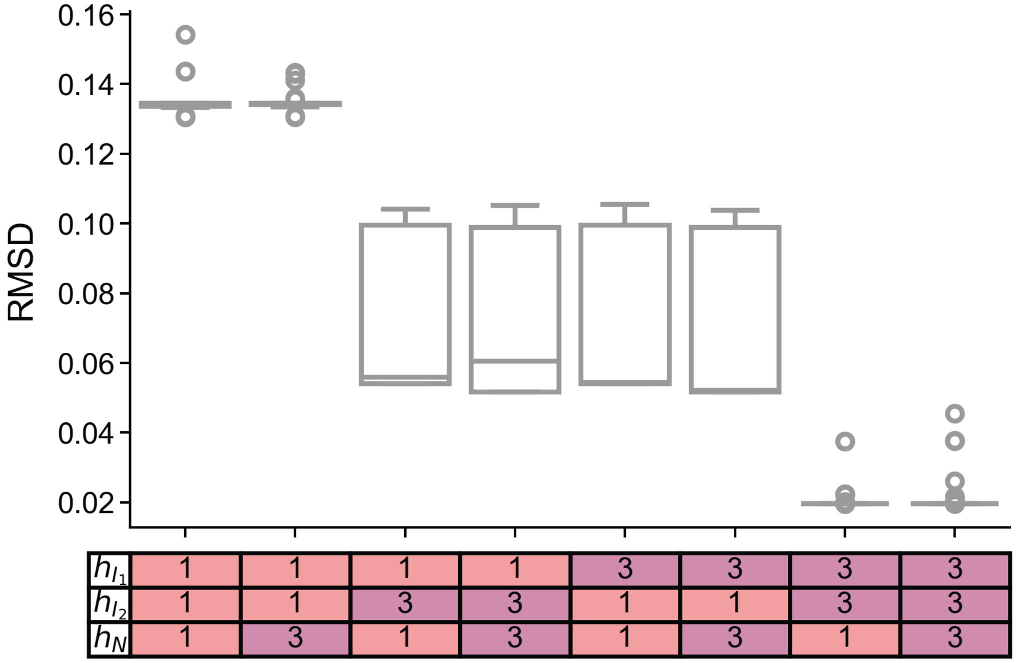


**Fig. S2.** Distributions of RMSD values for top 100 initially sampled parameters given multiple models of $h_{I_1}$, $h_{I_2}$, and $h_N$ for three-site model.

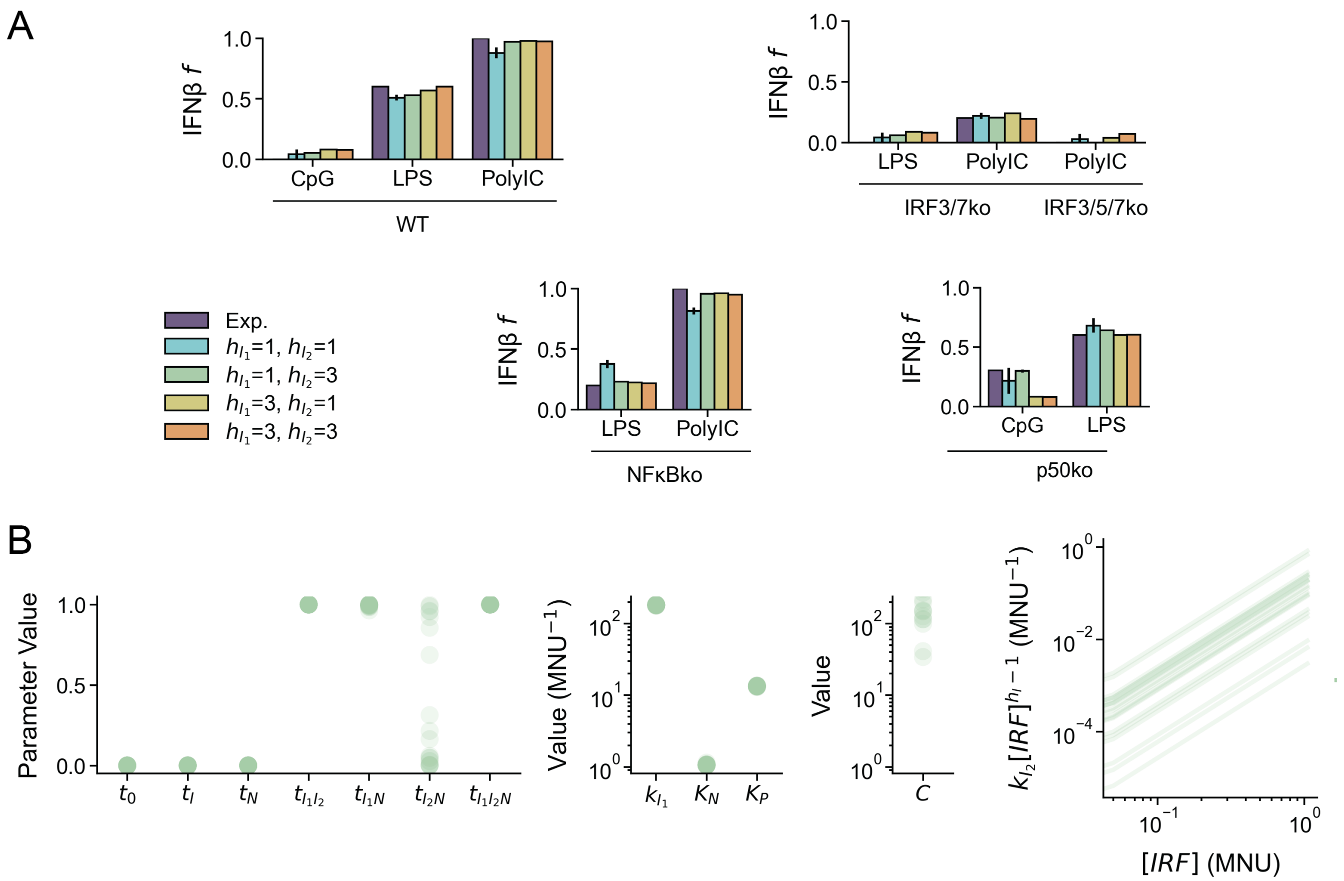


**Fig. S3.** (A) Experimental and mean predicted expression of IFNβ from best 20 optimized parameter sets for all data points using four models of $h_I$ for three-site model with p50:p50 binding competition and binding cooperativity. (B) Best 20 optimized $t$, $k_{I_1}$, $K_N$, and $K_P$ parameter values for all data points for 1&3 three-site model with p50:p50 binding competition and binding cooperativity.

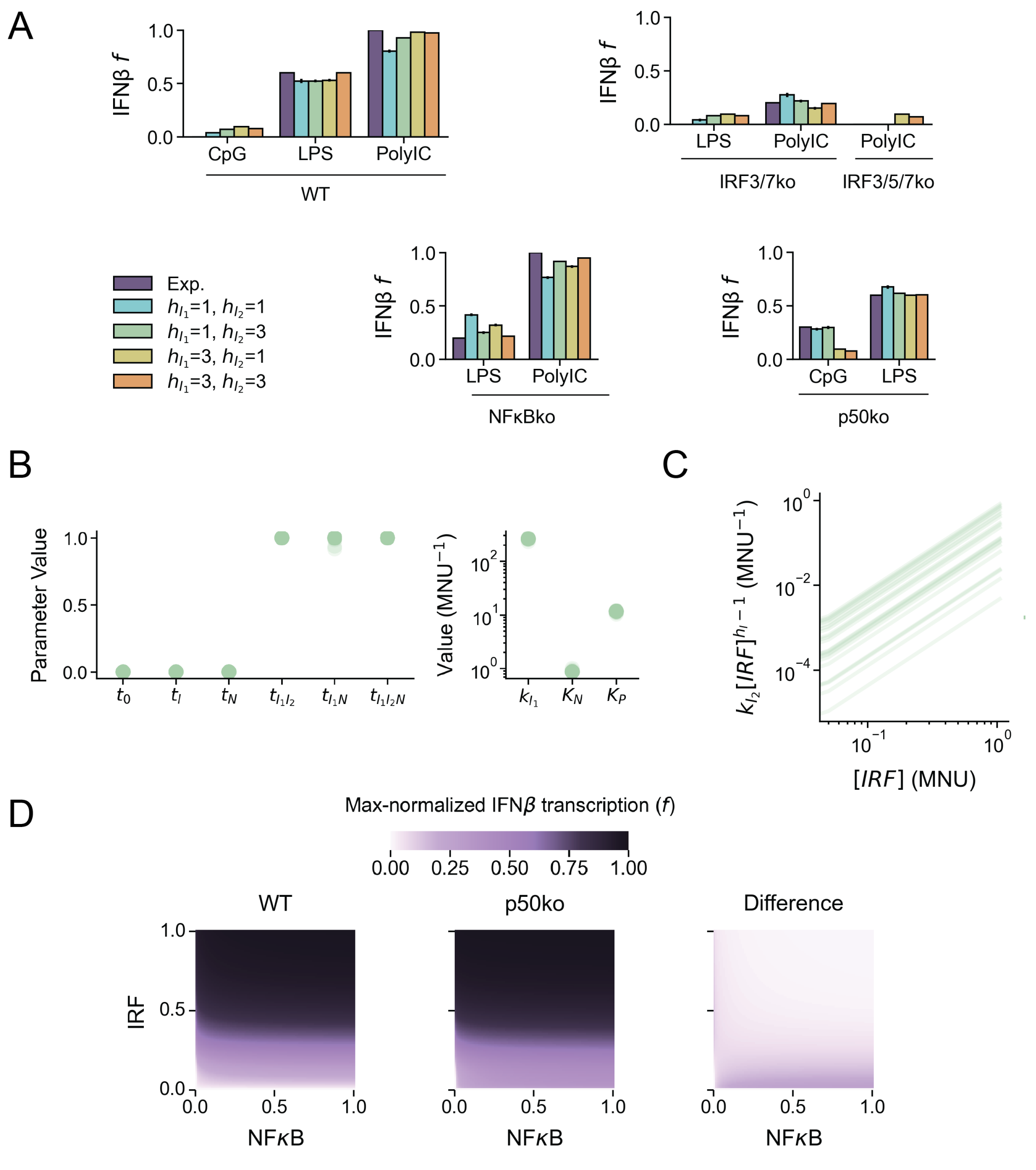


**Fig. S4.** (A) Experimental and mean predicted expression of IFNβ from best 20 optimized parameter sets for all data points using four models of $h_I$ for three-site model with p50:p50 binding competition without functional synergy between non-neighboring proteins (i.e., $t_{I_2N} = t_{I_2} + t_N$). (B) Best 20 optimized $t$, $k_{I_1}$, $K_N$, and $K_P$ parameter values for all data points for 1&3 three-site model with p50:p50 binding competition without functional synergy between non-neighboring proteins. (C) Binding affinity for IRE$_2$ as a function of $[IRF]$. (D) Total IFNβ transcription for different IRF and NFκB activities in WT p50 condition (left), p50ko condition (middle), and p50ko – WT (right).

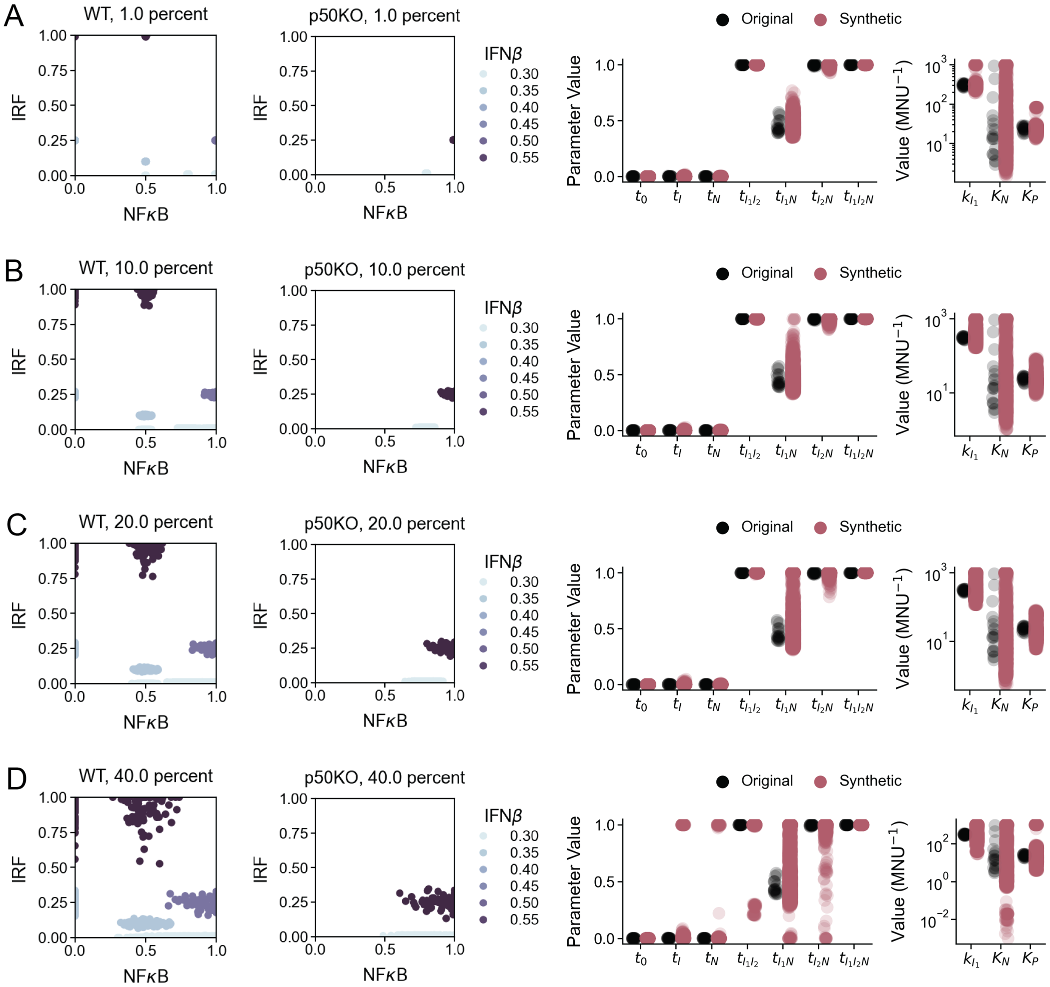


**Fig. S5.** (A-D) Synthetic data points in WT (left) and p50ko (middle) conditions and best 20 optimized $t$, $k_{I_1}$, $K_N$, and $K_P$ parameter values (right) for original and synthetic datasets with the following amount of error: (A) 1%, (B) 10%, (C) 20%, (D) 40%.

### References


1. T Kawai, S Akira, The role of pattern-recognition receptors in innate immunity: update on Toll-like receptors. *Nat. Immunol.* **11**, 373–384 (2010) Publisher: Nature Publishing Group.
2. S Akira, K Takeda, Toll-like receptor signalling. *Nat. Rev. Immunol.* **4**, 499–511 (2004) Number: 7 Publisher: Nature Publishing Group.
3. S Akira, S Uematsu, O Takeuchi, Pathogen recognition and innate immunity. *Cell* **124**, 783–801 (2006).
4. AL Blasius, B Beutler, Intracellular Toll-like Receptors. *Immunity* **32**, 305–315 (2010).
5. DN Rios, Ph.D. thesis (UC San Diego) (2014).
6. HM Lazear, et al., IRF-3, IRF-5, and IRF-7 Coordinately Regulate the Type I IFN Response in Myeloid Dendritic Cells Downstream of MAVS Signaling. *PLOS Pathog.* **9**, e1003118 (2013) Publisher: Public Library of Science.
7. M Sato, et al., Distinct and Essential Roles of Transcription Factors IRF-3 and IRF-7 in Response to Viruses for IFN-α/β Gene Induction. *Immunity* **13**, 539–548 (2000).
8. KL Peters, HL Smith, GR Stark, GC Sen, IRF-3-dependent, NFκB- and JNK-independent activation of the 561 and IFN-β genes in response to double-stranded RNA. *Proc. Natl. Acad. Sci.* **99**, 6322–6327 (2002).
9. S Sakaguchi, et al., Essential role of IRF-3 in lipopolysaccharide-induced interferon-β gene expression and endotoxin shock. *Biochem. Biophys. Res. Commun.* **306**, 860–866 (2003).
10. K Honda, et al., IRF-7 is the master regulator of type-I interferon-dependent immune responses. *Nature* **434**, 772–777 (2005).
11. X Wang, et al., Lack of Essential Role of NF-κB p50, RelA, and cRel Subunits in Virus-Induced Type 1 IFN Expression1. *The J. Immunol.* **178**, 6770–6776 (2007).
12. J Wang, et al., NF-κB RelA Subunit Is Crucial for Early IFN-β Expression and Resistance to RNA Virus Replication. *The J. Immunol.* **185**, 1720–1729 (2010).
13. X Wang, et al., Differential Requirement for the IKKβ/NF-κB Signaling Module in Regulating TLR- versus RLR-Induced Type 1 IFN Expression in Dendritic Cells. *The J. Immunol.* **193**, 2538–2545 (2014).
14. KA Ngo, et al., Dissecting the Regulatory Strategies of NF-κB RelA Target Genes in the Inflammatory Response Reveals Differential Transactivation Logics. *Cell Reports* **30**, 2758–2775.e6 (2020).
15. CS Cheng, et al., The Specificity of Innate Immune Responses Is Enforced by Repression of Interferon Response Elements by NF-κB p50. *Sci. Signal.* **4** (2011).
16. NE Buchler, U Gerland, T Hwa, On schemes of combinatorial transcription logic. *Proc. Natl. Acad. Sci.* **100**, 5136–5141 (2003) Publisher: Proceedings of the National Academy of Sciences.
17. MS Sherman, BA Cohen, Thermodynamic State Ensemble Models of cis-Regulation. *PLoS Comput. Biol.* **8**, e1002407 (2012).
18. P Virtanen, et al., SciPy 1.0: Fundamental Algorithms for Scientific Computing in Python. *Nat. Methods* **17**, 261–272 (2020).
